# Self-labeling of proteins with chemical fluorescent dyes in BY-2 cells and Arabidopsis seedlings

**DOI:** 10.1101/2020.03.09.983924

**Authors:** Ryu J. Iwatate, Akira Yoshinari, Noriyoshi Yagi, Marek Grzybowski, Hiroaki Ogasawara, Mako Kamiya, Toru Komatsu, Masayasu Taki, Shigehiro Yamaguchi, Wolf B. Frommer, Masayoshi Nakamura

## Abstract

Synthetic chemical fluorescent dyes are promising tools for many applications in biology. SNAP tagging provides a unique opportunity for labeling of specific proteins *in vivo* with synthetic dyes for studying for example endocytosis, or super-resolution microscopy. However, despite the potential, chemical dye tagging has not been used effectively in plants. A major drawback was the limited knowledge regarding cell wall and membrane permeability of synthetic dyes. Twenty-six out of 31 synthetic dyes were taken up into BY-2 cells, eight were not taken up and can thus serve for measuring endocytosis. Three of the dyes that were able to enter the cells, SNAP-tag ligands of diethylaminocoumarin, tetramethylrhodamine (TMR) and silicon-rhodamine (SiR) 647 were used to SNAP tag α-tubulin. Successful tagging was verified by live cell imaging and visualization of microtubules arrays in interphase and during mitosis. Fluorescence activation-coupled protein labeling (FAPL) with DRBG-488 was used to observe PIN2 endocytosis and delivery to the vacuole as well as preferential delivery of newly synthesized PIN2 to the newly forming cell plate during mitosis. Together the data demonstrate that specific self-labeling of proteins can be used effectively in plants to study a wide variety to cell biological processes.

## Introduction

Small-molecule fluorescent dyes have been used widely in biological research. Challenges in efficient use were however that they could not be targeted to specific cellular compartments or even proteins. Fluorescent proteins as genetically encoded dyes enabled efficient labeling of proteins in live or fixed material. In 2003, Kai Johnsson’s lab developed a tool for covalent labeling of fusion proteins with small molecules *in vivo, the* SNAP (O^6^-alkylguanine-DNA alkyltransferase) tag (Keppler et al., 2003). SNAP mediates covalent fusion of a dye of interest modified with a benzylguanine side chain to a specific target protein that carries the SNAP tag. Later, analogous tags such as Halo- and CLIP-tags were developed that enable simultaneous multicolor labeling of different proteins (Gautier et al., 2008; Los et al., 2005) react with O^2^-benzylcytosine, a cytosine conjugated to a dye, while the Halo-tag makes use of a mutated bacterial dehalogenase that creates stable bonds with any labeling chloroalkane ligand.

Over the past decade, many new dyes have been developed that open new possibilities such as ultra-photobleaching-resistant dyes for superresolution imaging with STED (Wang et al., 2017), endocytosis (Komatsu et al., 2011), and as sensors of biochemical parameters such as pH or metabolites (Asanuma et al., 2014; Jiang et al., 2019). Overall advantages of tagging of chemical dyes over other methods include: access to a wide range of fluorescent dyes, which typically have higher quantum yield and photostability compared to fluorescent proteins (FP), fluorescence is initiated when the dye is conjugated to the target protein, providing temporal control, as e.g. for DRBG-488 (Komatsu et al., 2011), and the tags are smaller relative to most FPs, thus likely less invasive. Tagging with different dyes has been widely used in the animal field for *in vivo* labeling (Yang et al., 2015). It has also successfully been deployed for *in vitro* studies of the role of the class II formin AtFH14 from *Arabidopsis* (Zhang et al., 2016). Surprisingly, only two reports have been published from the plant field. Both studies indicated that *in vivo* SNAP tagging may not be feasible in plants. One such example was the attempt to label apyrase. Labeling of fusion proteins *in vivo* with the cell-permeable, fluorescent substrates tetramethylrhodamine-Star and BG-505 gave high background, making the detection of AtAPY1-SNAP-specific fluorescence in Arabidopsis seedlings impossible (Schiller *et al.*, 2012). The tested dyes passed the cell wall and entered the cell, but even 14-h washing did not remove excess dye. A second study reported an attempt to use SNAP tagging for localizing the cytochrome *c* oxidase (COX) assembly factor HCC2 in mitochondria (Steinebrunner *et al.*, 2014). The authors were however unable to detect biotinylated HCC2-SNAP fusion proteins in mitochondria. Possible reasons could be that the commercially available dyes did not efficiently pass through the cell wall or the plant membranes are impermeable. Although unlikely, the conditions in the respective cellular compartments interfered with the labeling reaction.

Due to the large potential of small protein labeling for in cell biology, we systematically tested cell wall and plasma membrane retention of 31 different chemical dyes. We could classify dyes into three groups: dyes that enter the cytosol, dyes that enter in a pH-dependent fashion and dyes that can not be taken up by the cells. We subsequently used four different dyes to test SNAP tagging of different cargo by tagging microtubules and the auxin transporter PIN2. Here, we demonstrate self-labeling of microtubules with different dyes with different emission spectra in BY-2 cells and multi-color live cell imaging in the Arabidopsis seedling with genetically encoded fluorescent proteins. Localization and abundance of plasma membrane proteins such as transporters and receptors are dynamically controlled through endocytosis, recycling and vacuolar degradation in addition to *de novo* synthesis of the proteins (Luschnig and Vert, 2014; Yoshinari and Takano, 2017). Here we show that PIN2 undergoes clathrin-mediated endocytosis and subsequent vacuolar sorting and *de novo* synthesized PIN2 protein is preferentially transported to the cell plate rather than endocytosed/recycled PIN2 protein. The self-labeling of PIN2 shown here has advantages over alternative methods such as photoconversion of fluorescent proteins which is limited by the activation radius and artifacts that occur as a consequence of time lapse imaging. Taken together, our data show that SNAP tagging can be used for *in vivo* labeling in plants, thereby opening a wide range of applications to plant sciences.

## Results

### Uptake of synthetic dyes in BY-2 cells

We collected 32 different fluorescent dyes from commercial resources, colleagues and dyes developed at our institute (ITbM) (Table. 1). The list includes dyes that had been developed for SNAP tagging in animal cells, dyes for measuring endocytosis of receptors and dyes for super-resolution microscopy. To evaluate whether there is a fundamental problem for all dyes in passing through the cell wall, which dyes may be able to enter the cell, and to test whether certain dyes may be suitable for protein self-labeling, we systematically tested whether any of the 31 dyes (DRBG-488 is non-fluorescent before SNAP tagging) could enter BY-2 cells using confocal microscopy. Uptake experiments were performed in modified LS medium at pH 5.8 (Katsuta et al., 1990). The dyes could be classified into two categories based on the relative fluorescence intensity inside versus outside of the cell (Fig. 1; Supplementary Fig. 1, 2). 23 dyes were able to enter the cytoplasm of BY-2 cells within one minute, while 8 dyes did not show substantial uptake over the short exposure period (Fig. 1a, b; Supplementary Fig. 1, 2), e.g. Rhodamine Green (RG), Rhodamine 123 (a dye that accumulates in mitochondria of animal cells, which can be used for monitoring membrane potential), PREX710 (long wavelength dye for multicolor imaging) did not appear to be taken up to a measurable degree, and thus at least after short incubation times do not seem suitable for plant cell biology. Other dyes that might be suitable for monitoring endocytosis of membrane proteins such as DRBG-488 could not be tested since they carry a quenching group that is released when used for self-labeling. 2COOH RhP-M (pH sensitive dye for membrane labelling) was also not taken up efficiently but could therefore be suitable for measuring the pH during endocytosis in plant cells. While our survey was performed only for the one-minute time point, we also tested for two compounds whether uptake was possible when using longer incubation periods. While RG did not show any detectable uptake even after 30 min, the methylated variant 2MeRG, which showed some accumulation after 1 min, displayed multiphasic uptake kinetics and accumulated 40-fold within 30 min (Fig. 1c). Similar to 2MeRG, also diethylaminocoumarin, tetramethylrhodamine (TMR, protein labeling in immunohistochemistry) and 2MeSiR650 (single molecule measurements) accumulated in BY-2 cells (Fig. 1a). All the experiments above were performed at pH 5.8, a typical pH in the physiological range for the apoplasm. Since several dyes, e.g. hydroxymethyl-group bearing O-rhodamines (HM-rhodamines) alter their charge or change their structure in a pH dependent manner, we tested whether uptake was pH dependent. We found that HMRG showed increased accumulation at higher pH (Fig. 1d). Among four commercially available dyes frequently used for SNAP tagging in animals cells, SNAP-Surface Alexa Fluor 488 showed no accumulation in the cells but SNAP-Cell 430 as a diethylaminocoumarin derivative, SNAP-Cell TMR-star as a TMR derivative, and SNAP-Cell 647-SiR, a SiR650 derivative, showed low but detectable accumulation (Fig. 1e). In summary, permeable dyes suitable for SNAP tagging of imaging of cytoplasmic processes and cell impermeable dyes suitable for tagging membrane proteins at the cell surface were identified.

**Figure 1.**
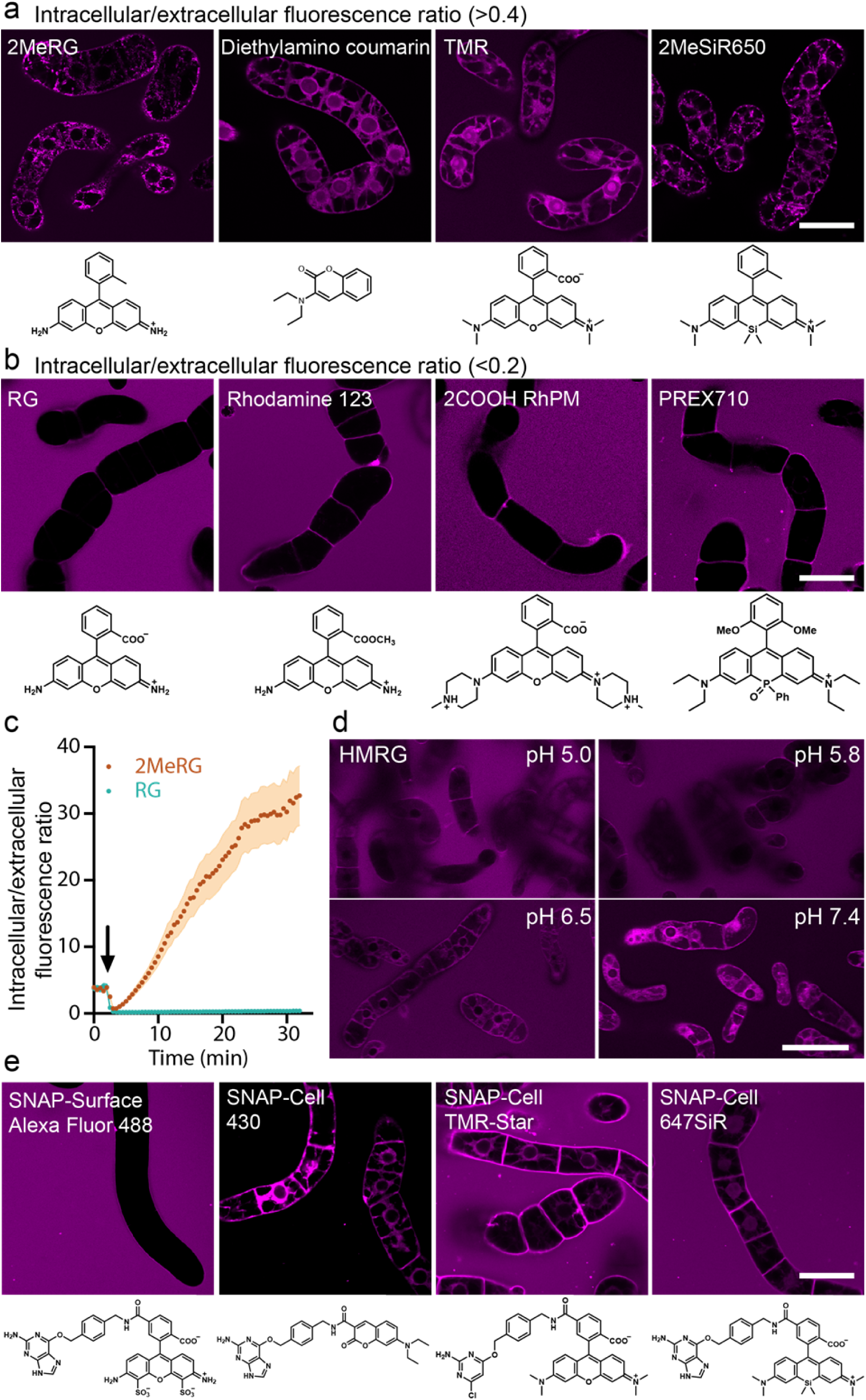
Representative images of BY-2 cells exposed to various fluorescent dyes and time course of the accumulation of fluorescent probes. **a**, Representative images of four dyes that enter BY-2 cells. Permeability was calculated by relative of fluorescence intensity inside / outside of the cell. Intra-/extracellular fluorescence intensity ratio **>**0.4 indicated uptake. **b**, Representative data for three dyes unable to enter BY-2 cells. Intra-/extracellular fluorescence intensity ratio <0.2. **c** Time-lapse analysis of 2MeRG and RG accumulation. Intra-/extracellular signal ratio of 2MeRG and RG. Plots and error bands represent mean (*n* = 6 cells) and SE, respectively. Arrow indicates time of dyes addition. Initial positive ratio is in absence of dye and due to autofluorescence. **d**, HMRG uptake at different pH. **e**, Confocal images of BY-2 cells incubated with SNAP-Surface Alexa Fluor 488, SNAP-Cell 430, SNAP-Cell TMR-Star and SNAP-Cell 647SiR for 1min. Images were taken with same settings at different time points. Scale bars: 50 µm. Experiments were repeated independently 3 times with comparable results.

**Figure 2.**
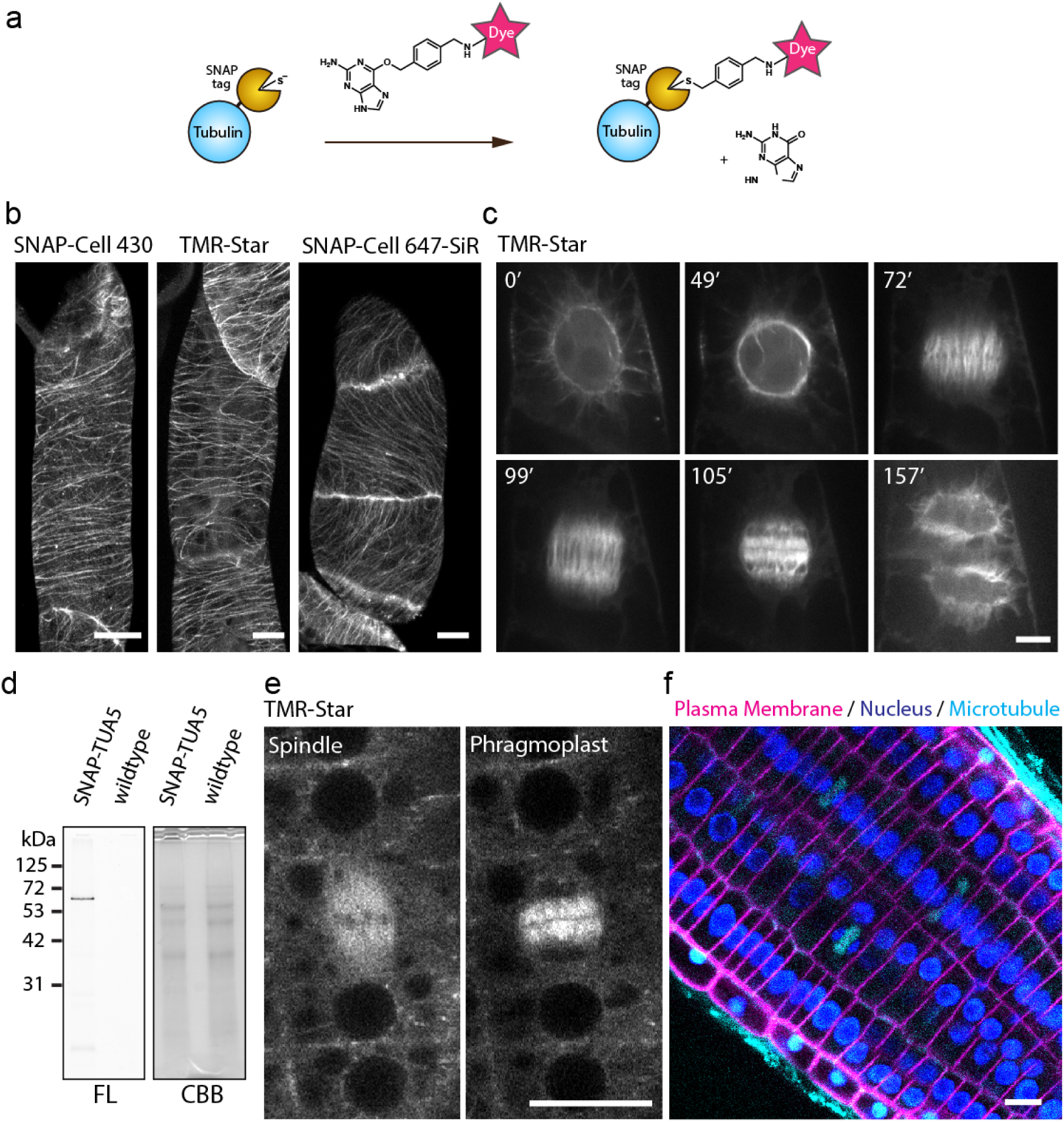
SNAP-tag enabled *in vivo* imaging of tubulin in BY-2 cells and Arabidopsis. **a**, Labeling mechanism of SNAP-tag. **b**, Cortical microtubules in SNAP-TUA5 expressing BY-2 cells with SNAP dyes denoted above images. Images show max intensity projection of confocal z-stack slices taken with 0.5 µm steps. **c**, Time-lapse imaging of mitotic microtubule dynamics TMR-star labeling of TUA5. Images taken every 30 sec, elapsed time (min) is shown. **d**, pUBQ10:SNAP-TUA5 and Col-0 (wildtype) seedlings were stained with 500 nM SNAP-Cell TMR-Star for 3h, lysed and analyzed by SDS-PAGE. Left panel: fluorescence; right panel Coomassie blue staining. **e**, Confocal images of mitotic cells in Arabidopsis root epidermis stained with TMR-Star. Spindles and phragmoplasts were observed. **f**, Root tip of Arabidopsis coexpressing p35S:YFP-LTI6b, p35:H2B-RFP, and pUBQ10:SNAP-TUA5. 3-day-old seedlings were incubated in 1/2MS containing 500 nM SNAP-Cell 647-SiR for 30 min. Scale bars: 10 µm. Experiments were repeated independently 3 times with comparable results.

### SNAP tagging of α-tubulin in BY-2 cells

While the experiments with BY-2 cells may indicate that there is no fundamental issue with dye permeability, other factors could be responsible for reported failure to deploy SNAP tagging in plant cells (Schiller et al., 2012; Steinebrunner et al., 2014). We therefore tested whether the commercially available dyes for self labeling SNAP-Cell 430, SNAP-Cell TMR-star and SNAP-Cell 647-SiR would be suitable for labeling the cytoskeleton of BY-2 cells. When BY-2 cells stably transformed with a construct pUBQ10:SNAP-α-tubulin (TUA5) were incubated with the any of the three dyes, efficient labeling of the microtubules was observed (Fig. 2). In interphase cells cortical microtubules were found to be highly ordered and oriented perpendicular to the growth axis (Fig. 2b). In mitotic cells microtubule first form a preprophase band in the preprophase, then generate the spindle during metaphase and subsequently transition to a concentrated cylinder, the phragmoplast during telophase (Fig. 2c; Supplementary Movie 1). The observed microtubule structures are highly similar to those seen when using fluorescent-protein tagged TUA5 (Lindeboom et al., 2013). Thus apparently, SNAP tagging with these dyes is feasible at least in tobacco BY-2 cell cultures.

### SNAP tagging of α-tubulin in Arabidopsis cells

The experiments with BY-2 cells may indicate that tag labeling is feasible in plants, however it is also conceivable that BY-2 cells as single cells or cell files were more accessible compared to cells in intact organs like plant roots. Alternatively, BY-2 cells as cell cultures might have particular features that allow passage of dyes through the cell wall and entry into the cytoplasm. We therefore tested whether the cytoskeleton could be labeled in a similar way in intact roots of Arabidopsis stably transformed with pUBQ10:SNAP-TUA5. SDS Page demonstrated that SNAP-Cell TMR-Star selectively labeled SNAP-TUA5 in protein extracts from Arabidopsis seedlings (Fig. 2d). Similar as in BY-2 cells, cytoskeletal structures such as spindle and phragmoplast were efficiently labeled with SNAP-Cell TMR-Star in Arabidopsis roots (Fig. 2e). One of the advantages of self-labeling with chemical dyes is the broad spectrum of dyes which enables multicolor in vivo imaging. Transgenic Arabidopsis plants expressing SNAP-TUA5 together with the plasma membrane marker (YFP-LTI6b) (Cutler et al., 2000) and the nuclear marker H2B-RFP (Federici et al., 2012). We chose the commercially available SNAP-Cell 647-SiR, a BG derivative of SiR650, a dye that had been shown to enter BY-2 cells (Fig. 1e), with a long wavelength emission, to label microtubules. After 30 min incubation with SNAP-Cell 647-SiR, microtubule arrays were detected together with the YFP-labeled plasma membrane and RFP-marked nuclei of Arabidopsis root tips (Fig. 2f; Supplementary Movie 2). Thus SNAP tagging appears to be an efficient and simple to use tool at least for self-labeling of abundant cytosolic proteins such as α-tubulin.

### SNAP tagging of the auxin transporter PIN2

Cell membrane-impermeable probes can be particularly suitable for studying endocytosis of membrane proteins such as transmembrane receptors and transporters (Komatsu et al., 2011; Geiger et al., 2013; Roed et al., 2014). Dyes that could not enter plant cells may be particularly suitable for studying endocytosis of membrane proteins in plants. The cell membrane-impermeable probe, DRBG-488 carries an intramolecular-FRET quencher group that is released when reacting with SNAP tag. Thus DRBG-488 specifically labels tagged proteins at the cell surface (Komatsu et al., 2011). As a result, DRBG-488 becomes fluorescent only after covalent linkage to a tagged protein of interest, providing high signal-to-background, e.g., for monitoring endocytosis. To test whether DRBG-488 can be used to observe endocytosis of a polarly localized plasma membrane transporter, we examined the SNAP-tagged auxin efflux carrier PIN2 (Adamowski and Friml, 2015). The SNAP tag was fused to the extracellular N-terminus of PIN2, which in addition carried mCherry in the central cytosolic loop to be able to monitor the production of PIN2. PIN2 was expressed under its native promoter (Fig. 3a, b). To validate that the two-tag fusion SNAP-PIN2-mCherry fusion did not negatively impact PIN2 activity in Arabidopsis, we complemented the *pin2/eir1-1* mutant and investigated agravitropic root growth (Roman et al., 1995). SNAP-PIN2-mCherry fusions restored normal gravitropic growth of *pin2/eir1-1* (Fig. 3c), demonstrating that SNAP-PIN2-mCherry was functional. Moreover, consistent with previous reports (Roman et al., 1995), confocal microcopy using RFP emission showed that SNAP-PIN2-mCherry localized preferentially in apical and basal domains of the plasma membrane of epidermis and cortex, respectively (Supplementary Fig. 3a, b). DRBG-488 effectively labeled SNAP-PIN2-mCherry within 30 min (Supplementary Fig. 4a, b). Both RFP and green fluorescence derived from the de-quenched DRBG-488 identified polar SNAP-PIN2-mCherry localization at the apical and basal membranes of epidermal and cortical cells, respectively (Supplementary Fig. 5). To monitor endocytic internalization of SNAP-PIN2-mCherry, a pulse-chase experiment using DRBG-488 was performed. In the cytoplasm, green fluorescence (DRBG-488) was only found in punctate structures, presumably corresponding to endosomes. 30 min after washout of the dye, green and red fluorescence (mCherry) colocalized in puncta in the cytoplasm (Fig. 3d). 210 min after washout DRBG-488-SNAP-PIN2 derived fluorescence was detected in vacuoles in addition to the punctate structures seen also earlier (Fig. 3d). Consistently, the ratio of fluorescence in the cytoplasm relative to the plasma membrane (C/P) gradually increased over time (Fig. 3e). These results indicate that DRBG-488 labeled SNAP-PIN2-mCherry at the cell surface first, then over a period of about an hour, SNAP-PIN2-mCherry is endocytosed and after about two hours is transported to the vacuole. To corroborate that PIN2 is endocytosed, pharmacological inhibition of endocytosis by ES9-17, an inhibitor of clathrin heavy chain, which is required for clathrin-mediated endocytosis, was performed (Dejonghe et al., 2019). ES9-17 reduced DRBG-488-derived fluorescence in both endosomes and vacuoles (Fig. 4a). The number of endosomes labeled by DRBG-488 was reduced significantly: 4.1 (± 0.2 S.E.) and 2.1 (± 0.1 S.E.) in mock and ES9-17 treatments, respectively (Fig. 4b). Similarly, The C/P ratio of DRBG-488 fluorescence was significantly lower: 0.21 (± 0.01 S.E.) and 0.103 (± 0.003 S.E.) in mock and ES9-17 treatments, respectively (Fig. 4c). The effect of ES9-17 on the C/P ratio of mCherry fluorescence was lower compared to DRBG-488 (Supplementary Fig. 6), due to continuous *de novo* synthesis of SNAP-PIN2-mCherry (mCherry positive, and DRBG-488-negative) in the cytoplasm. Taken together, DRBG-488 only labels SNAP-PIN2-mCherry localized at the cell surface and internalization of DRBG-488-labeled SNAP-PIN2-mCherry depends at least to a large extent on clathrin-mediated endocytosis.

**Figure 3.**
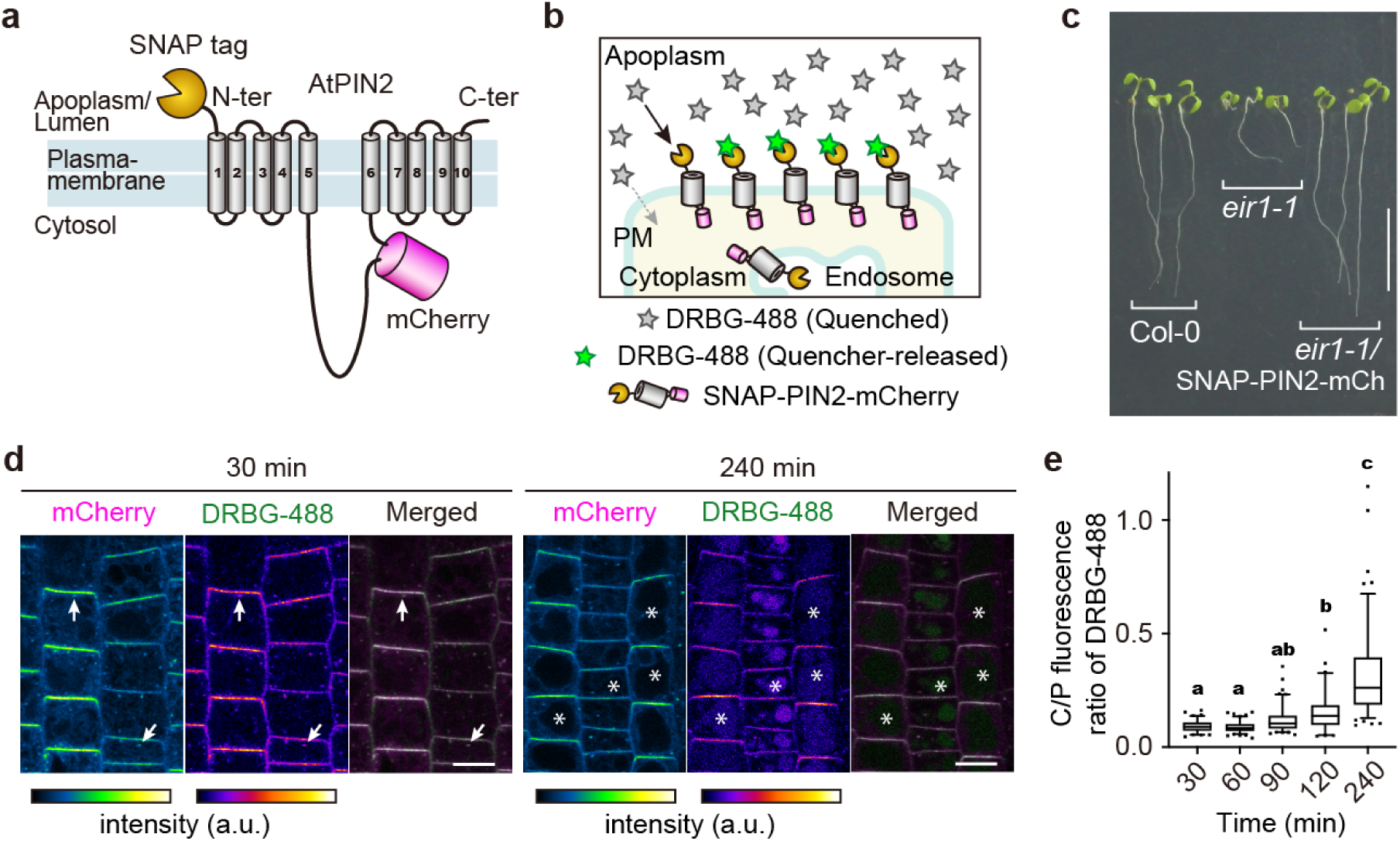
SNAP-tagging of PIN2 auxin transporter. **a**, Topological model of SNAP-PIN2-mCherry. SNAP-tag and mCherry are fused at the N-terminus and cytosolic loop of PIN2 protein, respectively. **b**, Conceptional illustration of labeling of SNAP-PIN2-mCherry by DRBG-488. DRBG-488 itself is impermeable and not fluorescent until reaction with SNAP tag. Once DRBG-488 binds to SNAP-tag, quencher group of DRBG-488 is released and becomes fluorescent. Fluorescent DRBG-488 reacts covalently with the SNAP-tag and is internalized with the chimeric SNAP-PIN2-mCherry. **c**, Phenotype of wild-type Col-0, *eir1-1, eir1-1* harboring *pPIN2:SNAP-PIN2-mCherry* grown for 7 days on solid media on vertical plates. Arrow indicates a root detached from the medium. Scale bar: 10 mm. **d**, pulse-chase analysis of DRBG-488 labeling. Plants were incubated with 200 nM DRBG-488 for 30 min followed by washout and 30- or 240-min incubation in liquid medium. Arrows indicate signals in plasma membrane and intracellular punctate structure. Asterisks indicate vacuoles. Scale bars: 10 µm. **e**, Cytoplasm/plasma membrane (C/P) fluorescence ratio of DRBG-488. Box plots are centered at the data median and mark from the 25th to the 75th percentile. Dots indicate outliers. Different letters above the plots represent significant differences among means [*P* < 0.005 by one-way analysis of variance (ANOVA) with Tukey’s post-hoc test. Exact *P* values are described in Supplementary Data]. *N* = 86, 104, 104, 62, and 118 cells from three roots. Experiments were repeated independently 3 times.

**Figure 4.**
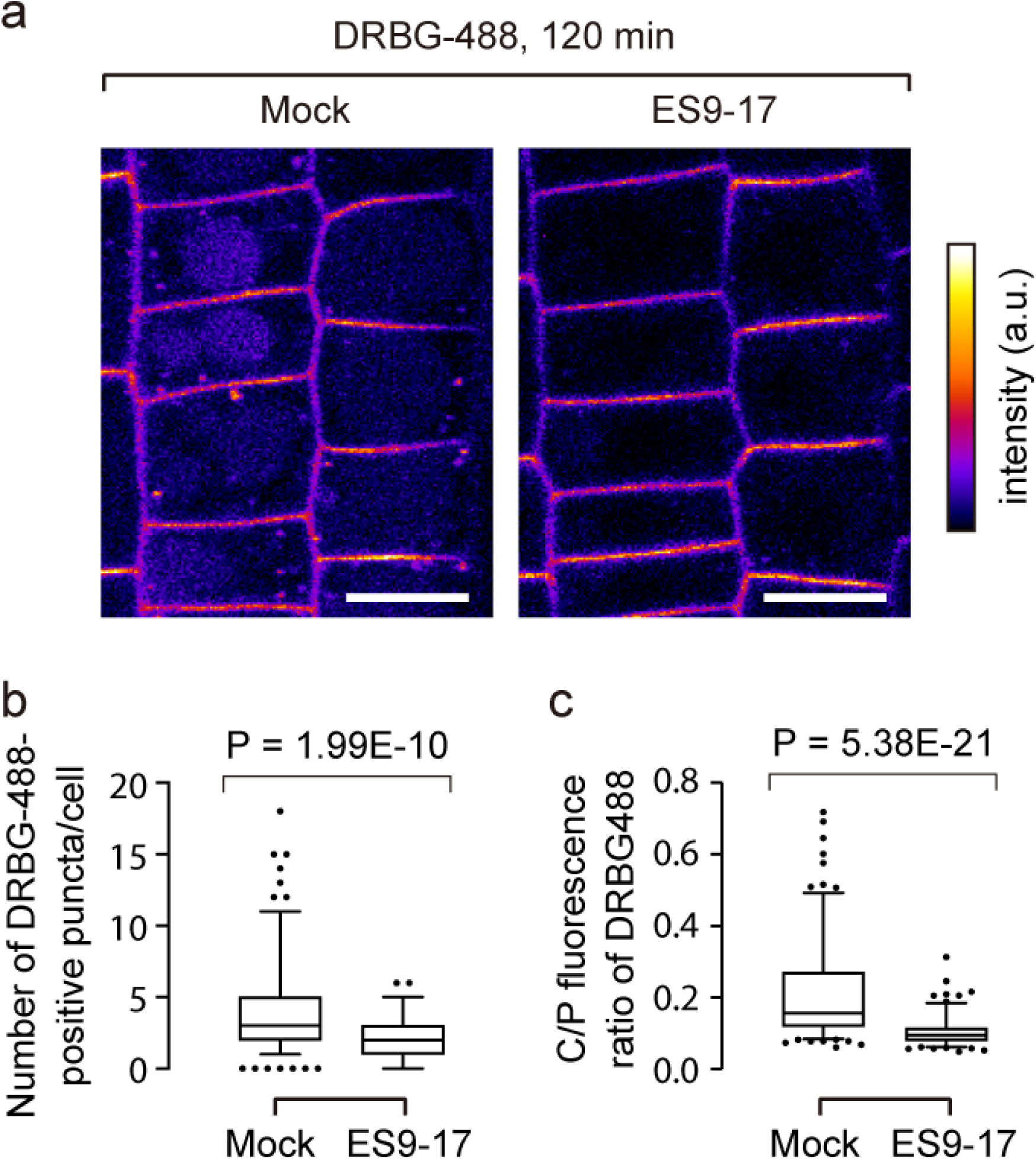
Evidence for clathrin-mediated endocytosis of PIN2. **a**, DRBG-488-labeling of SNAP-PIN2-mCherry with DMSO (mock) or 30 µM ES9-17 for 120 min. Scale bars: 10 µm. **b**, Number of DRBG-488-positive endosomes per cell in the presence/absence of ES9-17. Significance of difference was determined by two-tailed Student’s *t*-test. *n* = 177 (DMSO) and 111 (ES9-17) cells from 5 independent roots. **c**, C/P fluorescence ratio of DRBG-488 in the presence/absence of ES9-17. Box plots are centered at the data median and mark from the 25th to the 75th percentile. Dots indicate outliers. Significance of difference was determined by a two-tailed Student’s *t*-test. *n* = 177 (DMSO) and 158 (ES9-17) cells from 5 roots. Experiments were repeated independently 3 times.

**Figure 5.**
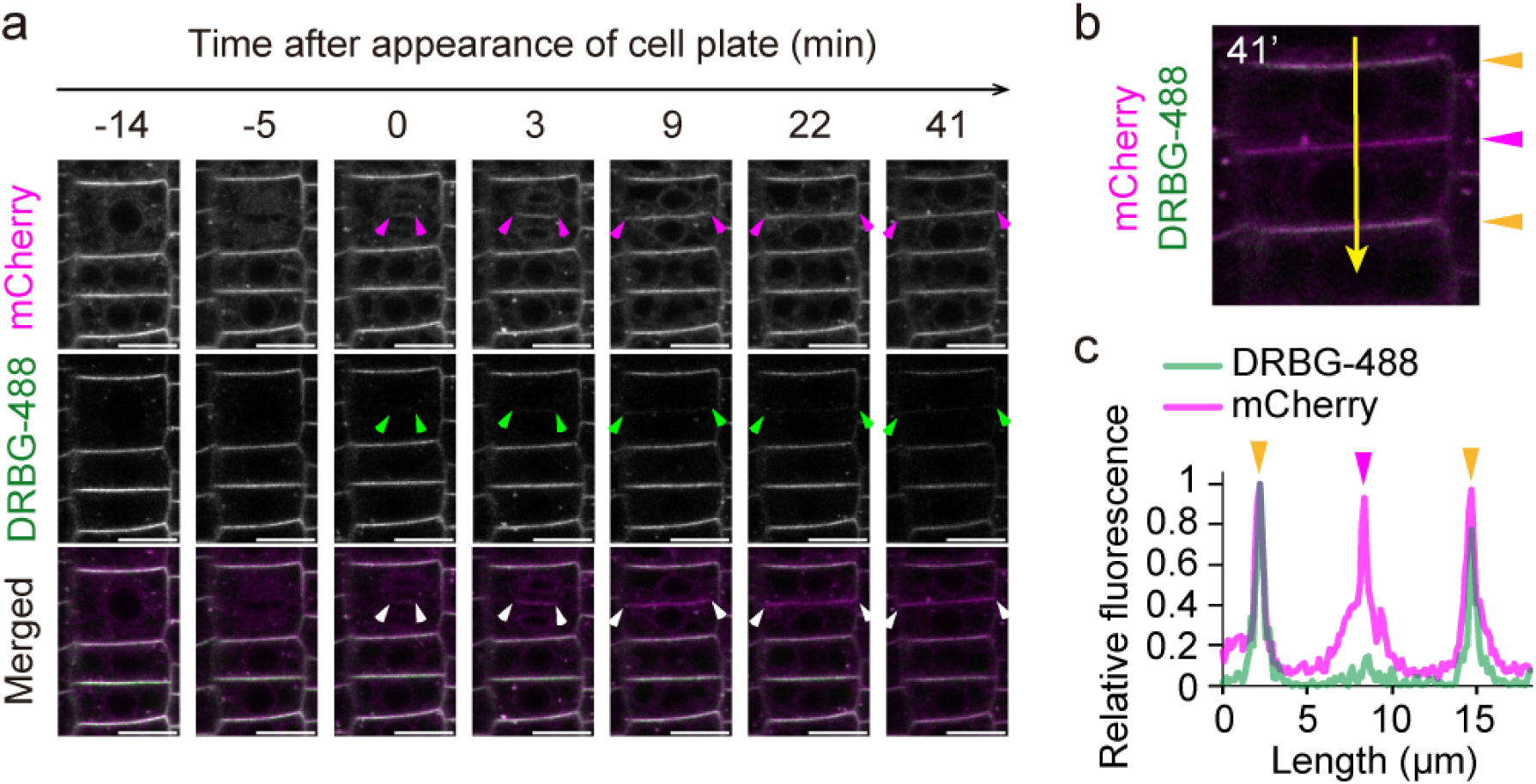
Preferential delivery of newly produced PIN2 to the newly forming cell plate. **a**, Time-lapse imaging of PIN2 a dividing cell. Arrowheads indicate edges of the cell plate and connection sites between newly-formed plasma membrane and the lateral plasma membrane. Scale bars: 10 µm. **b**, Quantification of fluorescence intensities of DRBG-488 and mCherry along the line in the cell at 41 min after appearance of cell plate. Orange and pink arrowheads indicate pre-existing and newly-formed plasma membrane, respectively. **c**, Scan of relative fluorescence of DRBG-488 and mCherry in images from panel b. Colored arrowheads on peaks correspond scan position in b. Experiments were repeated independently three times.

It had previously, been reported that *de novo* synthesized PIN2 is preferentially transported to the cell plate during cytokinesis, followed by re-distribution of PIN2 in a polar manner (Mravec et al., 2011; Glanc et al., 2018). We thus hypothesized that the DRBG-488-free fraction of SNAP-PIN2-mCherry, which represents *de novo* secreted PIN2 will accumulated at higher levels at the cell plate relative to the DRBG-488-labeled fraction, which is subject to endocytosis. We thus determined the ratio of mCherry/DRBG-488 fluorescence in different parts of the plasma membrane of dividing epidermal cells in the Arabidopsis root over time. Time-lapse imaging after labeling with 200 nM DRBG-488 for 90 min showed that while the ratio of DRBG-488 to mCherry-labeled SNAP-PIN2-mCherry was approx. 1 in the apical part of the cells, the newly formed cell plate had a ratio of DRBG-488 to mCherry-labeled SNAP-PIN2-mCherry of >10 (Fig. 5). These data are consistent with the hypothesis that endocytosis of PIN2 from the apical domains is not contributing to a large extent to the PIN2 amount present at the cell plate which is mainly derived from *de novo* synthesis. In summary, the reaction of a membrane-impermeable self-quenched dye with the SNAP tag on the extracellular N-terminus of a polytopic transporter in combination with the fusion of a fluorescent protein enabled us to monitor endocytosis, intracellular trafficking and delivery of newly synthesized protein to the cell plate.

## Discussion

Self-labeling of proteins *in vivo* is a tool that has several advantages of the use of fusions with fluorescent proteins. Unfortunately it appeared that this extensive tool set with many different dyes was not applicable to plant cells for unknown reasons. Given the large potential of the technology and the wide range of advantages and possibilities that chemical dyes provide, we reevaluated the utility in plants in a systematic way. A systematic screen of 31 dyes allowed is to classify the dyes into cell permeable and impermeable compounds. We subsequently demonstrated SNAP tagging of microtubules in BY-2 cells using three structurally very different dyes. We also showed multicolor labeling with SNAP tagging in combination with FPs. Using a quenched dye that is unable to enter cells on its own to observe endocytosis of the SNAP-tagged auxin transporter PIN2.

### Synthetic dyes

The analysis of a total of 32 chemical dyes shows that some dyes are able to enter BY-2 cells within less than a minute. Other dyes enter more slowly and for some, no substantial uptake was observed. In addition, the permeability of the plasma membrane of BY-2 cells was pH dependent. For some dyes also uptake into roots of intact Arabidopsis plants was observed, indicating that the survey in BY-2 provides a good way to determine whether such dyes can or can not be taken up by plant cells. Notably commercially available dyes used for SNAP tagging that can enter animal cells also pass the cell wall and plasma membrane of plant cells. Dyes that enter the cells can in principle be used for self-labeling of cytosolic and nuclear proteins and possibly proteins in subcompartments such as vacuole, plastids and mitochondria. Dyes that cannot enter the cells can in principle be used to label proteins present at the cell surface such as transporters and receptors. One potential caveat is that for self-labeling, the dyes need to be modified by addition of a benzylguanine or analogous reagent, which may affect permeability. Thus for dyes that have not been tested in their SNAP form, permeability must be determined. We could not measure the permeability of DRBG488, a dye that carries both the benzylguanine and a quencher that largely eliminates fluorescence of unreacted reagent. It only becomes fluorescent once it is SNAP-tagged to a protein at the cell surface. We could however confirm that DRBG488 labeled the auxin transporter PIN2 specifically at the cell surface. Taken together, the survey (together with the results from SNAP tagging show that a wide range of dyes with unique properties can in principle be used by self-labeling of proteins inside cells or at the cell surface.

For self-labeling of cytosolic proteins, SNAP-Cell 430, SNAP-Cell TMR-Star and SNAP-Cell 647SiR were used to visualize microtubule arrays in BY-2 cells. By making use of the long wavelength excitation and emission maximum, SNAP-Cell 647SiR was also used to perform three-color live-cell imaging in Arabidopsis seedling. The near-infrared (NIR) fluorescent dyes (650-900 nm) are useful for fluorescence imaging of mammalian cells and tissues, since the tissue transparency is high and within the NIR optical window autofluorescence are dramatically reduced in mammalian tissues (Weissleder and Ntziachristos, 2003). Although plants have phytochromes with absorption maxima at 660 and 730 nm (Li et al., 2011), as demonstrated here NIR dyes with a self-labeling technique are deemed suitable for multicolor NIR fluorescence imaging in plant cells. In addition to SiR-based dyes, PREX670 and BcPR705 have long wavelength spectra and across the plasma membrane, which also permit simultaneous use with commonly used genetically fluorescent protein probes. The high photostability of PB430 and its relatively long fluorescence lifetime (**Τ** = 10.6 ns) makes it suitable for stimulated emission depletion (STED) microscopy imaging (Wang et al., 2017). The result of permeability indicates that synthesis of SNAP-PB430 dye would be beneficial for super resolution imaging of cytosolic proteins. Dyes such as 2COOH RhP-M could be used as a pH sensor in endocytosed compartments since it was unable to enter the cytoplasm.

### Factors contributing to the difference in dye permeability

In principle there are three major routes by which dyes can be taken up by a cell: (i) passive diffusion across the lipid membrane, (ii) recognition and uptake via endogenous transporters or (iii) endocytosis. Overton’s rule stated that transport across a membrane correlates with its lipid solubility, thus charge and hydrophobicity (Overton, 1895). Overton’s rule has been challenged (Grime et al., 2008). Transporters have been identified for compounds that were previously thought to move passively across lipid bilayers, e.g. water uses aquaporins (Preston et al., 1992), ammonium uptake is mediated by AMTs (Ninnemann et al., 1994), and fatty acids are taken up by fatty acid transporters (Schaffer and Lodish, 1994). Moreover, many transporters have been show to transport more than just one substrate, e.g. through the peptide transporter PepT1 (Brandsch, 2013). Charge, shape and hydrophobicity are also important aspects of recognition by a transporter. Thus at present we can not differentiate how the dyes shown here to be cell permeable move across the plasma membrane. Possible indications may come from a careful analysis of the dyes for which we observed pH dependent uptake. Presumably, multiple factors may affect the cell permeability of compounds, such as structure, charge, 3D conformation, and hydrophobicity. Hydroxymethyl-group bearing O-rhodamines (HM-rhodamines) such as HMRG, HMDiMeR and HMTMR undergo intramolecular spirocyclization equilibrium between open form and spirocyclized ‘closed form’ with a p*K*_cycl_ (pH at which absorbance of the compound decreases to half maximal absorbance as a result of spirocyclization) around the physiological pH of mammalian cells at 7.4. This equilibrium has been utilized for measuring pH (Kenmoku et al., 2007) or to increase the activation rate of fluorescent probes (Kamiya et al., 2011; Sakabe et al., 2013; Matsuzaki et al., 2016). (Supplementary Fig. 7). One min after dye addition, the time window over which we measured the transport of the dyes in the screening, HMRG, with no substitution on *N*, entered the cells comparatively slowly, whereas HMDiMeR and HMTMR were readily taken in. This could be due to increased hydrophobicity of the HM-rhodamines by addition of methyl groups on *N*, or an increase in p*K*_cycl_ with the addition of methyl groups. An increase in hydrophobicity was confirmed by HPLC and values of CLogP, showing that HMDiMeR and HMTMR are increasing hydrophobic as more methyl groups are added to HMRG (Supplementary Fig. 7b). Increase in p*K*_cycl_ means that the ratio of open form decreases at a given pH. If there is a difference in permeability between open and closed forms, compounds with more permeable forms at the assay pH would be taken up more readily. Further studies will be required before we can draw any conclusions on the key factors that drive cell membrane permeability for these dyes.

### Dynamic analysis of endocytosis and intracellular trafficking of PIN2 by SNAP-tagging

FAPL with DRBG-488 had successfully been used to monitor endocytosis of the EGF receptor in COS7 and MDCK animal cell cultures (Komatsu et al., 2011). Also in plants, endocytosis and intracellular trafficking play important roles in maintenance of membrane protein abundance and subcellular localization (Luschnig and Vert, 2014). Here we utilized a cell-impermeable fluorescent probe DRBG-488 that is de-quenched upon reaction with the SNAP tag for labeling of SNAP-PIN2-mCherry at the cell surface. Our data intimate that PIN2 is internalized by clathrin-mediated endocytosis (Fig. 4). This is consistent with the conclusion from previous studies (Dhonukshe et al., 2007; Kitakura et al., 2011; Adamowski et al., 2018; Glanc et al., 2018). These studies used photoconvertible fluorescent proteins, pharmacological experiments with brefeldin A (BFA) and genetic evidence. BFA is widely used to examine endocytic rate in plant researches. BFA inhibits some subsets of ADP ribosylation factor guanine nucleotide exchange factors (ARF-GEFs) such as GNOM. GNOM, however, is localized to both the plasma membrane and the Golgi but not in the *trans*-Golgi network/early endosomes (TGN/EEs) in normal conditions and BFA induces dissociation of GNOM from Golgi apparatus and plasma membrane and accumulation in the TGN/EE (Naramoto et al., 2010, 2014). Consequently, BFA disturbs not only endosomal recycling indirectly but also endocytosis of membrane proteins (Naramoto et al., 2010, 2014). Furthermore, BFA alters morphology of the Golgi apparatus and induces absorption of the Golgi cisternae into endoplasmic reticulum in aerial tissues of Arabidopsis, implying that there are unknown molecular targets other than the ARF-GEFs (Robinson et al., 2008). Thus, BFA treatment compromises GNOM-dependent endocytosis and natural intracellular trafficking, making it difficult to draw reliable conclusions on endocytosis. Photoconvertible fluorescent proteins (FP) such as Dendra2 (Jásik et al., 2013; Glanc et al., 2018) are also widely used to monitor endocytosis of proteins. However, photoconvertible proteins potentially undergo undesirable photoconversion during time-lapse imaging in addition to difficulty of photoconversion of restricted regions of interest. Moreover, photoconversion occurs in a sphere that has a radius of several micrometers thus activating also proteins inside the cell. For application with cell surface proteins, self labeling thus has two major advantages: it targets only proteins at the cell surface and permits high resolution imaging that is otherwise limited by the undesirable photoconversion of FPs. An advantage of the photoconvertible FPs is the ability to mark proteins specifically in a subset of the plasma membrane, which is not possible with a simple SNAP tagging approach. Furthermore, we confirmed by pulse-chase observation of DRBG-488-labeled SNAP-PIN2-mCherry that newly-formed PIN2 protein is transported to the cell plate rather than endocytosed/recycled PIN2 protein (Fig. 5). This result is consistent with observations from a previous report in which a photoconvertible FP was instead used (Glanc et al., 2018). Together we demonstrate SNAP-tagging prides a new way for dynamic analysis of endocytosis and intracellular trafficking of functional membrane proteins such as PIN2.

Taken together, over 30 chemical dyes were classified in BY-2 cell-permeable and impermeable compounds. Self-labeling of soluble proteins with chemical dyes that are taken up into plant cells via a SNAP tag can be efficiently used to fluorescently tag the microtubule cytoskeleton to observe cytoskeletal dynamics *in vivo*. Self-labeling expands e.g. the options for multicolor labeling and enables tagging novel dyes e.g. pH sensors or compounds suitable for superresolution microscopy such MitoPB Yellow, PhoxBright 430, or PREX 670 (Wang et al., 2017, 2017; Grzybowski et al., 2018). Self-labeling of soluble proteins with chemical dyes via a SNAP tag can be efficiently used to fluorescently tag membrane proteins at the cell surface, in particular for FAPL. Here we used DRBG-488, that is plant cell impermeable also in Arabidopsis and to tag PIN2 at the cell surface and to monitor endocytosis. We demonstrate that SNAP-tagged PIN2 accumulates at newly forming cell plates to higher levels than mCherry labeled PIN2, providing support for the hypothesis that newly synthesized PIN2 is delivered to the cell plate rather than being endocytosed.

## Materials And Methods

### Sources and synthesis of dyes

Dyes were either obtained from commercial sources, from the Yamaguchi (ITbM, Nagoya University) or the Urano lab (The University of Tokyo). 6-[[4-(aminomethyl) phenyl] methoxy]-9*H*-purin-2-amine (BG-NH_2_) was prepared according to previous reports (Keppler et al., 2004). TMR (tetramethylrhodamine) and 2COOH RhP-M were synthesized here (for full details see Supplementary Methods).

#### DNA constructs

The pUBQ10:SNAP-TUA5 binary vectors were generated using Gateway technology (Invitrogen) by following the Multisite Gateway Technology three-fragment vector construction kit protocol. UBQ10pro-p4p1r and t35S-p2rp3 constructs were kindly provided by Jelmer Lindeboom and David Ehrhardt (Carnegie Science, Stanford). The SNAP-tag sequence was amplified using pSNAPf vector (#N9183, NEB) and inserted into the *Asc*I and *Kpn*I sites after UBQ10 promoter in the UBQ10pro-p4p1r entry vector. The open reading frame of AtTUA5 (At5g19780) was cloned into pDONR/zeo. Three entry clones, UBQ10pro-p4p1r, TUA5-pDONR/zeo and t35S-p2rp3 were transferred into the binary vector pHm43GW (Karimi et al., 2005) by LR Clonase II Plus enzyme (Invitrogen). The plasmid map of pSTA5 is shown in Supplementary Fig. 8. For the construction of SNAP-PIN2-mCherry, the genomic sequence of *AtPIN2* (At5g57090; −2784 bp from the start codon to +688 bp from the stop codon) was amplified from *Arabidopsis* genomic DNA by PCR and integrated into pGEM-T Easy vector (Promega). First, mCherry was introduced into the PIN2 construct by inverse PCR followed by an In-Fusion reaction (Clontech). The SNAP-tag with an N-terminal secretion signal peptide sequence (MKTNLFLFLIFSLLLSLSSAEF; the signal sequence likely improves import of the tag into the ER, since constructs lacking the signal sequence did not show mCherry fluorescence) was synthesized by IDT gBlocks® Gene Fragments service (Supplementary Tables S1 and S2). The SNAP-tag was introduced into the PIN2-mCherry construct by inverse PCR followed using the In-Fusion reaction. The resulting construct was digested by *Not* I and integrated into pBIN40 binary vector by ligation (Miyashima et al., 2011). The resulting binary vector was introduced into *Agrobacterium tumefaciens* strain GV3101:MP90 and used to transform Arabidopsis and tobacco BY-2 cells. The plasmid map of pAY230 is shown in Supplementary Fig. 8.

#### Plant cell cultures and SNAP tag labeling with fluorescent dyes

BY-2 cells were cultured in modified Linsmaier and Skoog (LS) medium (Katsuta et al., 1990) buffered with 0.05% (w/v) MES pH 5.8. Cells were subcultured once a week by adding 1.5 ml (wild type cells) or 3 ml (SNAP-TUA5 expressing cells) of the cell suspension into 80 ml fresh medium. Cultures were maintained in the dark at 25 °C under rapid shaking (120 rpm). To generate transgenic BY-2 cell lines expressing SNAP-TUA5, Agrobacterium-mediated transformation was carried out basically following published methods (http://2010.igem.org/Team:Nevada/BY-2_(NT1)Transformation_Protocol). Transformants were selected in LS medium containing 0.8% (w/v) agar, 40 µg ml^-1^ hygromycin, and 200 µg ml^-1^ carbenicillin. Resistant calli was transferred to new selection media, then left 3 weeks before initiating liquid culture in modified LS medium. To label SNAP-TUA5 with chemical dyes, 1-day or 2-day old SNAP-TUA5 expressing BY-2 cells was used. 500 µl subcultured cells were transferred into 1.5 ml tubes, and media were replaced with SNAP-dye containing medium. For SNAP-Cell TMR-star (#S9105S, NEB) and SNAP-Cell 430 (#S9109S, NEB), solutions were prepared by diluting 1 mM dye stock solution in 100% DMSO to a final concentration 500 nM. For SNAP-Cell SiR647 (#S9102S, NEB) was also prepared from 1 mM dye stock solution in DMSO to a final concentration 1 µM, but to solve the dye into the media the final DMSO concentration was adjusted to 1% (v/v). Incubation times varied between 5-60 min depending on the dye and experiments; 5 or 10 mins for SNAP-Cell TMR-star, 30 mins for SNAP-Cell 430, and 60 mins for SNAP-Cell 647-SiR. After incubation with SNAP-dyes, cells were washed with 3% (w/v) sucrose several times, then resuspended in fresh LS medium. For time lapse imaging of microtubules, cells were attached to poly-L-lysine coated glass-based dishes.

#### Plant Material and Treatments

For multicolor labeling we generated Arabidopsis lines expressing p35S:H2B-RFP (nuclei)(Federici et al., 2012), p35S:YFP-LTI6b (plasma membrane)(Cutler et al., 2000) and pUBQ10:SNAP-TUA5 (here) by crossing. Partially heterozygous F2 was used for imaging. Plants were grown on 0.5x Murashige and Skoog (MS) media with 1 % (w/v) sucrose and 1 % (w/v) agar or modified MGRL media (Takano et al., 2005) containing 1 % (w/v) sucrose and 1.5 % (w/v) gellan gum (Wako Pure Chemicals, Osaka, Japan) and 30 µM boric acid. Seeds were surface-sterilized with 70% (v/v) ethanol and sown on media plates. To synchronize germination, plates were placed in the dark at 4 ºC for 2 days. Plants were grown vertically in a 16-h/8-h light/dark cycle (40 μE m^−2^ s^−1^) for 4 days for imaging, or 7 days for growth analysis. For fluorometric analysis on SDS-PAGE (sodium dodecyl sulfate-polyacrylamide gel electrophoresis), 4-day-old seedlings were incubated in 0.5x MS medium containing 500 nM SNAP-Cell TMR-star for 3 hours. Cell extracts were lysed in 50 mM Tris-HCl at pH 7.5, 150 mM NaCl, 1% Triton X-100 and 2 mM phenylmethylsulfonyl fluoride and Complete Protease Inhibitor cocktail (Roche Diagnostics). 2 µg of the extracted proteins were separated by SDS-PAGE gel and stained with CBB R-250. Fluorescent images were captured by Typhoon FLA 9500 (GE Healthcare). For microtubule imaging, plants were soaked in 0.5x MS medium containing 500 nM SNAP-dye. For DRBG-488 staining, all experiments were performed in liquid culture. Seedlings were soaked in MGRL solution (modified MGRL liquid medium without both sucrose and gellan gum) containing 200 nM of DRBG-488 under light condition (∼120 μE m^−2^ s^−1^). Notably, cells in the root tip and elongation zone showed reliable and specific labeling of PIN2, while cells in the calyptra, cells in older zones of the root and dead cells became also fluorescent, likely as a consequence of unspecific removal of the quencher from DRBG-488 in these cells. All reactions and analyses were performed at 23 – 25 ºC. For pulse-chase labeling by DRBG-488, plants were first treated with 200 nM DRBG-488 for 30 min, then residual dye was washed out by adding fresh MGRL solution. Plants were subsequently incubated in MGRL in the light (∼120 μE m^−2^ s^−1^) for up to 210 min with recordings every 30 min.

#### Confocal Imaging

Fluorescence images were captured using a Leica Application Suite Advanced Fluorescence (LAS-AF) instrument with a TCS SP8 gSTED and a 20 x objective (0.75 NA HC PL APO CS2, Leica), 63 x oil-immersion objective (1.40 NA HC PL APO CS2, Leica), or a 100 x oil-immersion objective (1.40 NA HC PL APO CS2, Leica). The light sources were a white-light laser or an Argon laser or a Diode laser. The excitation wavelength was set according to the maximum wavelength of dye absorption, 458 nm for SNAP-Cell 430, 488 nm for DRBG-488, 552 nm for SNAP-Cell TMR-star, 561 nm for mCherry, and 645 nm for SNAP-Cell SiR647. Detection with Leica HyD (GaAsP hybrid detection system) was set to 50 nm range according to the maximum wavelength of dye fluorescence, 463-650 nm for SNAP-Cell 430, 500–550 nm for DRBG-488, 557-764 nm for SNAP-Cell TMR-star, 570–650 nm for mCherry, and 648-749 nm for SNAP-Cell SiR647. Images were taken at 400 Hz, a picture size of 1024×1024 pixels and 1.0 AU. A time-gate pulse of 0.3 ns–6.0 ns was used for detection DRBG-488, SNAP-Cell TMR-star, and SNAP-Cell SiR647 to reduce background signals. All dyes were prepared in DMSO (Wako Pure Chemicals, Osaka, Japan) as a stock solution. Concentration of dyes was adjusted based on their absorbance. For screening of synthetic dyes using BY-2 cells (Fig.1 and Supplementary Fig. 1), 100 µl of cells in LS media at pH 5.8 were loaded into 8-well chamber slides (Iwaki), and 1 min after 100 µl 2µM dye solution were added into the well fluorescent images were taken using Leica Application Suite Advanced Fluorescence (LAS-AF) instrument with a TCS SP8 gSTED and a 20 x objective (0.75 NA HC PL APO CS2, Leica). The excitation wavelength was set according to the maximum wavelength of dye absorption and 50 nm range of wavelength according to the maximum wavelength of fluorescent dyes were detected with Leica HyD (GaAsP hybrid detection system). To change the pH of the assay, 500 µl of BY-2 cells in LS media were left to sediment for 5 min, then the supernatant was removed and replaced with 500 µl of buffer with a different pH. This procedure was repeated three times before cells were imaged as above. For time-course analysis of staining of BY-2 cells with 2MeRG and RG (Fig. 1C), 100 µl of cells in LS media at pH 5.8 were loaded into 8-well chamber slide (Iwaki) and fluorescent images were taken every 30 seconds using Leica Application Suite Advanced Fluorescence (LAS-AF) instrument with a TCS SP8 gSTED and a 20 x objective (0.75 NA HC PL APO CS2, Leica). 100 µl 2µM dye solution were added into the well between 120 sec and 150 sec. The excitation wavelength was set to 498 nm and 499 nm for 2MeRG and RG, respectively and the detections with Leica HyD (GaAsP hybrid detection system) were set to 50 nm range of 510-560 nm. For time lapse imaging of SNAP-dye labeled microtubules in BY-2 cells (Fig. 2C), an inverted microscope Eclipse Ti2 (Nikon) equipped with a spinning disk confocal unit CSU-10 (Yokogawa) and a sCMOS camera PRIME 95B (Photometrics) was used. SNAP-Cell TMR-star was excited with a 561 nm laser, and emission signals were collected with 605/64 nm filter (Semrock). Images were taken with a 60 x oil-immersion lens (CFI Plan Apo VC 1.40 NA, Nikon).

#### Quantitative analyses of PIN2 dynamics

PIN2 dynamics was analyzed in epidermal cells of meristematic and transition zones of primary roots. Image quantification was performed with Fiji/ImageJ software (ver. 2.0.0-rc-69/1.52p) (Steinebrunner et al., 2014). The number of endosomes were counted using an ImageJ macro, the SiCE spot detector with manual correction (Bayle et al., 2017). Manual correction was used by ignoring puncta that overlapped with the plasma membrane, and by manually assigning puncta that localized too close to plasma membrane to be detected automatically. Finally the number of puncta per cell was counted. For calculation of the C/P (cytoplasm to plasma membrane) fluorescence ratio, the mean fluorescence value in the apical plasma membrane domain detected by a segmented line tool with 3-pixel-thickness was defined as fluorescence in the plasma membrane and mean fluorescence value in the entire cytoplasm selected by a polygonal selection tool was assigned to the cytoplasm. Scatter plots were generated and statistical analyses were performed with Prism8 (GraphPad Software, CA).

#### Pharmacological inhibition of endocytosis

As described before, all DRBG-488 labeling experiments were performed in liquid culture. ES9-17 (Carbosynth, CA; cat# BE17203) was dissolved in DMSO at a concentration of 50 mM and stored in glass vial at −30 ºC. Plants were incubated with 30 µM ES9-17 or equivalent volume of DMSO (mock) diluted in MGRL solution for 30 min followed by incubation in MGRL solution containing 200 nM DRBG-488 and either 30 µM ES9-17 or equivalent volume of DMSO for 120 min with supplemental light next to the microcope (∼120 μE m^−2^ s^−1^) until imaging was started (base level at microscope without supplemental lighting: ∼10 μE m^−2^ s^−1^). The light intensity was chosen since mCherry fluorescence in the vacuole is apparently pH dependent and thus lower intensity is observed at higher light intensities (Tamura et al., 2003)

#### Imaging of cytokinesis in root epidermal cells of *Arabidopsis*

To label most of SNAP-PIN2-mCherry existing in the cells, plants were treated with 200 nM DRBG-488 for 90 min. Plants were then transferred into glass-bottom chambers and overlaid with an MGRL soft-gel medium pad [modified MGRL medium containing 0.8% (w/v) gellan gum without sucrose]. Root epidermal cells were observed by inverted confocal microscopy. During the time course experiment under the microscope, plants were exposed only to low light conditions, not taking the laser irradiation into account during imaging (∼1 μE m^−2^ s^−1^).

## Acknowledgements

We would like to thank Dr. Shunsuke Oishi (ITbM, Nagoya Unievrsity) for HPLC and Dr. Yoshikatsu Sato (ITbM Live Imaging Center, Nagoya University) for microscopes. We are grateful to Prof. Miyo Morita (National Institute of Basic Biology) for providing *eir1-1* seeds, Dr. David W. Ehrhardt (Carnegie Institution for Science) for YFP-LTI6b and H2B-RFP seeds, and Dr. Keitaro Umezawa (Tokyo Metropolitan Institute of Gerontology), Dr. Wen Piao, Dr. Kenjiro Hanaoka (the University of Tokyo) and Dr. Chenguang Wang (ITbM, Nagoya University) for providing their compounds. This research was supported by the Human Frontier Science Program to M.N., JSPS grant (18KK0195) to M.N. and A.Y., and the Alexander von Humboldt Foundation to W.B.F. ITbM is supported by World Premier International Research Center Initiative (WPI), Japan.

## Author Contributions

R.J.I., M.N and W.B.F conceived the projects. R.J.I., A.Y., N.Y., and M.N. performed the experiments. R.J.I., H.O., M.G., M.K., T.K., and M.T. synthesized, prepared and analyzed fluorescent dyes. M.N., W.B.F., R.J.I., A.K., and N.Y. analyzed the data and wrote the manuscript. All authors discussed the results and commented on the manuscript.

